# Improving Metagenomics Classification with Kmask: Entropy-Based Masking of Low-Complexity Regions

**DOI:** 10.64898/2026.09.22.753623

**Authors:** Yuchen Ge, Edward Li, Harun Mustafa, Ales Varabyou, Steven L. Salzberg

## Abstract

Accurate taxonomic classification in metagenomics is often compromised by low-complexity sequences, which lead to chance matches that in turn cause sequences to be misclassified. Here we present Kmask, an entropy-based masking tool implemented for use either standalone or as part of Kraken [1,2] database construction, which replaces low-entropy regions with Ns. Using a sliding window size aligned with Kraken’s default k-mer length and parameters optimized across 12 control bacterial genomes spanning a broad range of GC content, Kmask efficiently removes low-complexity sequences while retaining high-complexity regions. To benchmark performance, we applied Kmask to a newly constructed database, Microbial2025, that contains over 71,000 bacterial, archaeal, viral, and fungal genomes, and we then classified human reads against both masked and unmasked versions of the database using KrakenUniq [2]. We found that Kmask substantially reduced misclassifications, driving down the false positive rate to 5.78% from 7.52%. Notably, Kmask performed comparably to an SDUST-masked [3] database, achieving a similar false positive rate (5.78% vs. 5.17%) while masking out fewer bases (1.33% vs. 1.85%). We also tested Kmask on a database of human cancer sequences, where we found that it eliminated many false positives caused by low-complexity matches between bacterial genomes and human DNA. These results demonstrate that Kmask is an effective method for masking low-complexity sequences in large microbial databases, thus improving the accuracy of metagenomic classification.

## Introduction

In genomics, low-complexity sequences are DNA strings containing simple repeats such as long monomer runs, dinucleotide repeats, or other non-random sequences. Because unrelated genomes can contain similar low-complexity sequences, such regions can produce spurious matches that are not due to shared evolutionary history. A variety of sequence complexity measures have been proposed to distinguish informative from repetitive genomic regions, but they differ substantially in formulation and behavior. Classical approaches such as SDUST [3] quantify overrepresented short nucleotide patterns (e.g., triplets) and flag regions with biased composition, making those methods effective for identifying simple repeats but relatively insensitive to more subtle sequence structure. In contrast, measures based on Shannon entropy [4] can treat a sequence window as a distribution over k-mers and quantify its information content. These methods provide a principled, information-theoretic interpretation of complexity and can be tuned via k-mer length to match downstream applications, though they are sensitive to window size and sampling noise. Compression-based metrics, such as those derived from Lempel-Ziv complexity [5], estimate how well a sequence can be algorithmically compressed, capturing longer-range redundancy but often at higher computational cost. Other approaches, including linguistic complexity and repeat-based heuristics, explicitly measure the diversity or periodicity of substrings within a window. Collectively, these methods trade sensitivity to different types of structure with the statistical assumptions of downstream analyses, highlighting the importance of selecting a complexity definition that is appropriate for the intended application.

Among existing methods, the SDUST algorithm [3] (often referred to as “dust”), which defines complexity based on 3-mer frequencies, is widely used for detecting local low-complexity regions. However, its computational cost scales with both window size and sequence length, and its scoring function increases quadratically with sequence length, biasing the method toward classifying longer regions as low complexity. This limitation can become problematic when analyzing long tandem or satellite repeats. Recently, an extension of SDUST called longdust [6] was developed, which introduces a statistical model of k-mer count distributions and achieves near-linear time complexity, enabling efficient detection of long low-complexity regions. Notably, longdust improves sensitivity to extended repetitive regions that are often missed by SDUST, although it relies on approximate algorithms and still faces practical constraints on window size for very long repeats.

Here, we introduce Kmask, a masking method based on the standard Shannon entropy definition applied to k-mers. Rather than introducing a new statistical model, Kmask focuses on practical optimization of key parameters, including the choice of unit length and entropy threshold score, to better align with the behavior of k-mer-based metagenomics classifiers such as Kraken. This design enables effective identification of low-complexity regions that disproportionately contribute to ambiguous k-mer matches, while preserving informative sequence content. Kmask is implemented with an efficient sliding-window strategy that achieves near-linear time complexity of O(L), where L is the genome length, making it suitable for genome-scale database construction and large metagenomics workflows.

## Results

The first part of our study focuses on characterizing low-complexity sequences and optimizing Kmask’s parameters. We benchmarked different unit sizes for entropy calculation using a set of representative bacterial genomes spanning a broad range of GC content. Based on these results, we selected parameters that best balanced the goal of masking out low-complexity regions versus preserving high-complexity ones. We then applied these parameters across all genomes in a large database that we created, called Microbial2025 (see **Table 1**), to compute entropy scores for every k-mer, followed by empirical determination of the optimal entropy threshold that achieved effective masking with minimal loss of informative sequence content.

**Table 1.**
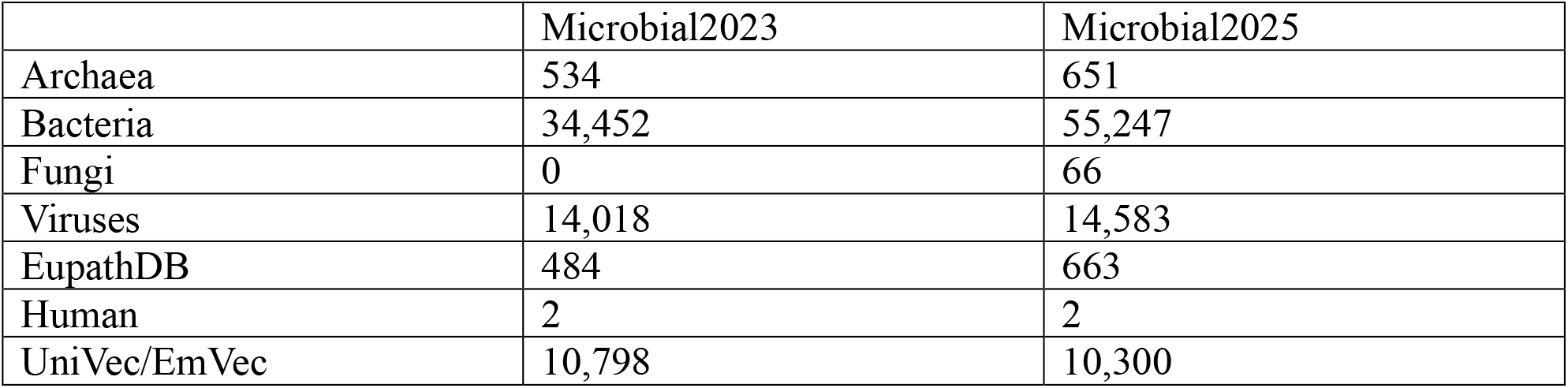
Genome counts by taxonomic category for the Microbial2023 and Microbial2025 reference databases.

|  | Microbial2023 | Microbial2025 |
| --- | --- | --- |
| Archaea | 534 | 651 |
| Bacteria | 34,452 | 55,247 |
| Fungi | 0 | 66 |
| Viruses | 14,018 | 14,583 |
| EupathDB | 484 | 663 |
| Human | 2 | 2 |
| UniVec/EmVec | 10,798 | 10,300 |

The second part of our study addresses the masking and construction of the Microbial2025 database. We built a separate KrakenUniq [2] database comprising genomes from common laboratory and environmental contaminants, including host-derived sequences, cloning vectors, and microbial taxa frequently misclassified in metagenomic analyses. Each genome in the Microbial2025 collection was then screened against this contamination database using KrakenUniq, and all 31-mers matching target contaminant sequences were masked in the corresponding genomes. This procedure removed shared or misassigned k-mers, yielding a cleaner and more reliable reference database.

Last, we evaluated the performance of the masked Microbial2025 database on a set of suspicious low-count species detected in human tumor samples from the TCGA project [7], previously published by our group. Comparisons against existing reference databases showed that the masked Microbial2025 database achieves improved precision in pathogen detection relative to its unfiltered counterpart.

### Choosing the best entropy unit length (*l*) for Kmask

We constructed 84 small KrakenUniq databases, each containing the same 12 bacterial genomes but using different combinations of *l* and *s* to mask the bacteria, where *l* is the entropy unit length and *s* is the entropy score threshold (Supplementary **Figure S1** and **Table S1**). The bacteria were chosen to represent a wide range of GC-content, from 27-73% (see **Table 2**). For testing, we created a set of “pseudo-reads” by taking every 31-bp sequence from the CHM13 human genome and treating it as a read (see Methods). By construction, any pseudo-read that was classified as one of the 12 bacteria would be a false positive.

**Table 2.** Bacterial genomes used in experiments to adjust key parameters of Kmask, sorted by GC content.

|  |  |  |  |
| --- | --- | --- | --- |
| Scientific name | Taxonomy ID | Assembly accession | GC content (%) |
| *Fusobacterium nucleatum* subsp. *nucleatum* ATCC 25586 | 190304 | GCF_003019295.1 | 27 |
| *Staphylococcus epidermidis* | 1282 | GCF_006094375.1 | 32 |
| *Helicobacter pylori* | 210 | GCF_025998455.1 | 38.5 |
| *Streptococcus pneumoniae* | 1313 | GCF_001457635.1 | 39.5 |
| *Bacillus subtilis* subsp. *subtilis* str. 168 | 224308 | GCF_000009045.1 | 43.5 |
| *Escherichia coli* str. K-12 substr. MG1655 | 511145 | GCF_000005845.2 | 51 |
| *Salmonella enterica* subsp. *enterica* serovar Typhimurium str. LT2 | 99287 | GCF_000006945.2 | 52 |
| *Neisseria meningitidis* | 487 | GCF_022869645.1 | 52 |
| *Pseudomonas putida* NBRC 14164 | 1211579 | GCF_000412675.1 | 62.5 |
| *Mycobacterium tuberculosis* H37Rv | 83332 | GCF_000195955.2 | 65.5 |
| *Streptomyces clavuligerus* | 1901 | GCF_005519465.1 | 72.5 |
| *Frankia alni* ACN14a | 326424 | GCF_000058485.1 | 73 |

For each run, we generated a report capturing the percentage of pseudo-reads classified as bacterial. To calculate the false positive rate (FPR), we used the number of pseudo-reads classified against the unmasked database as the baseline (*s* = 0). As the entropy threshold *s* increased, the number of classified reads decreased monotonically, until the database was fully masked and no reads were classified. The FPR at each unit length and threshold was therefore calculated as 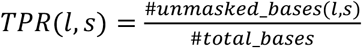

To quantify the true positive rate (TPR), we calculated the percentage of unmasked bases in the bacterial genomes, reasoning that these unmasked regions represent the sequence available for correct taxonomic classification. Under the assumption that a metagenomic sequencing dataset comprised reads that were sampled uniformly from these twelve bacteria, and that these reads were classified using a database containing the same twelve genomes after masking, the TPR can be approximated as the fraction of each genome’s bases remaining unmasked. Therefore 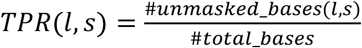.

As shown in **Figure 1**, unit sizes of *l* = 3 and *l* = 4 both achieved comparably high AUC values, substantially exceeding those observed for *l* = 1 and *l* = 2. The ROC curves for *l* = 3 and *l* = 4 were nearly indistinguishable across the tested threshold range. This overlap indicates that increasing the unit size beyond *l* = 3 provides negligible additional discriminative power, and that AUC alone is therefore insufficient to justify selecting *l* = 4 over *l* = 3.

**Figure 1.**
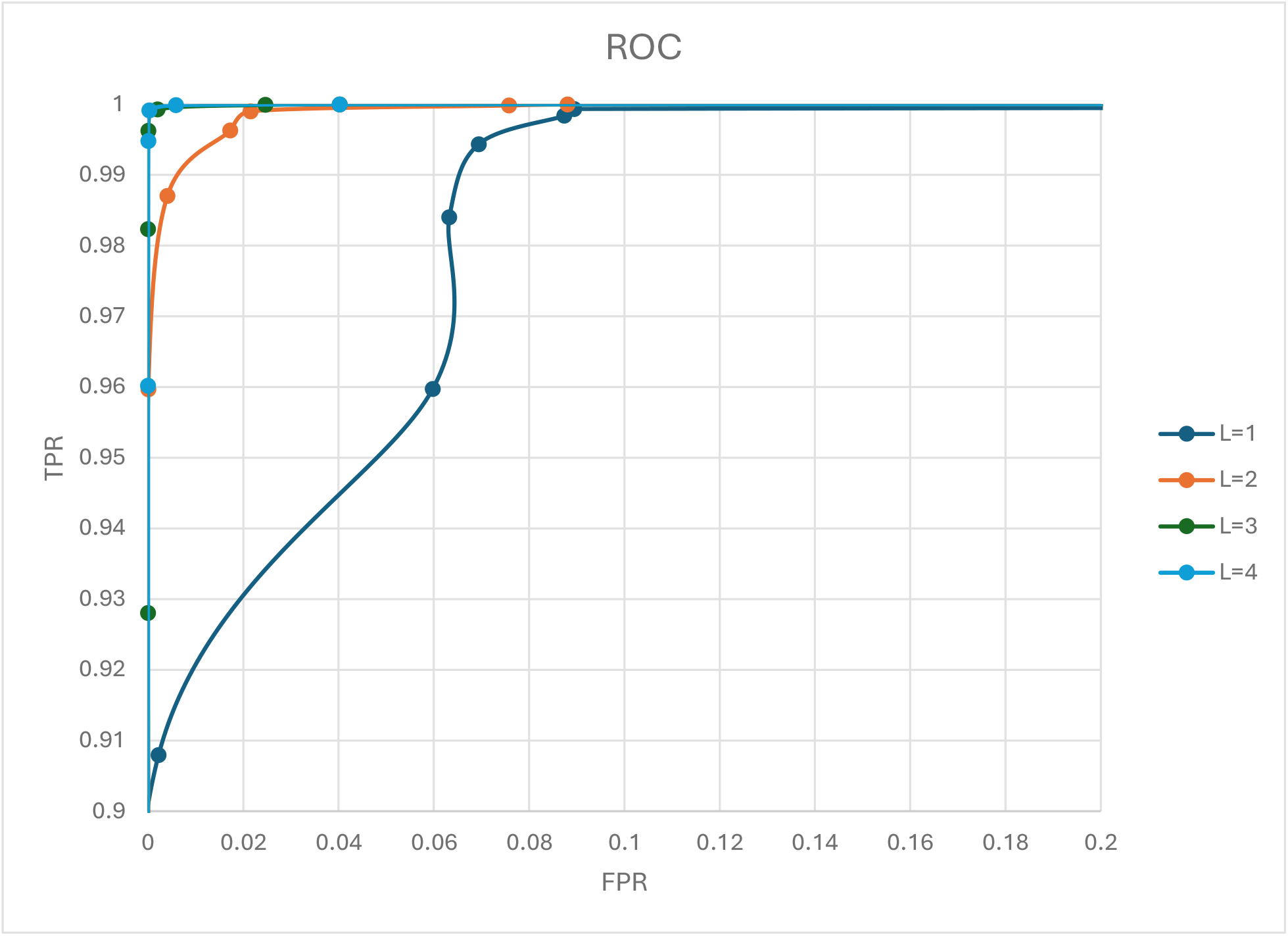
Receiver operating characteristic (ROC) curves evaluating Kmask performance across different unit sizes (*l*) for entropy-based masking. For each unit size (*l* = 1, 2, 3, 4), the true positive rate (TPR) and false positive rate (FPR) were calculated across a range of entropy thresholds using 31-bp pseudo-reads derived from the CHM13 human genome, and queried against 12 bacterial genomes masked at each threshold.

To further differentiate between these two choices, we considered how effectively each unit size utilizes its theoretical entropy range given the fixed k-mer length of 31-bp used throughout this study. The theoretical maximum entropy for a given unit size L is determined by the number of possible distinct *l*-mers (4^*l*^), yielding a maximum of log_2_ 4^*l*^ = 2*l* bits. For *l* = 3, this corresponds to a theoretical maximum entropy of 6 bits. However, because entropy is computed over the *l*-mers contained within each 31-bp k-mer, the number of overlapping L-mer positions available is limited to *k* ™ *l* + 1. For *l* = 3, this yields at most 29 unique 3-mer positions per k-mer, capping the practically achievable entropy at log2(29) ≈ 4.86 bits, or approximately 81% of the theoretical maximum.

By comparison, for *l* = 4 the theoretical maximum entropy rises to 8 bits, but the number of available 4-mer positions within a 31-bp k-mer is only 28, limiting the practically achievable entropy to log2(28) ≈ 4.81 bits, which is just 60% of the theoretical maximum. This means the entropy scores computed at *l* = 3 more fully occupy the dynamic range available to them, whereas at *l* = 4 a large portion of the theoretical entropy scale (6–8 bits) is effectively unreachable given the fixed k-mer length, compressing the usable resolution for threshold selection.

Taken together, these results indicate that *l* = 3 offers the more efficient and better-calibrated choice for entropy-based masking at *k* = 31: it achieves classification performance nearly identical to *l* = 4 while making fuller use of its theoretical entropy range. We therefore selected *l* = 3 as the optimal unit size for all downstream masking analyses.

### Construction of the Microbial2025 database

We built a comprehensive microbial reference database, Microbial2025, to serve as a curated and contamination-filtered resource for metagenomic classification and pathogen detection. The database consists of a total of 71,210 genomes available from the NCBI RefSeq database [8] as of December 31, 2025, spanning all major domains of life. Specifically, Microbial2025 includes 55,247 bacterial, 651 archaeal, 66 fungal, and 14,583 viral genomes, together with 663 eukaryotic pathogen genomes obtained from EuPathDB release 68 [9] which includes many widely studied protozoan and helminth pathogens (see Supplementary **Table S2**). Nearly all of the genomes had complete or near-complete assemblies, with the exception of the EuPathDB collection, which predominantly contains draft assemblies. **Table 1** summaries the main contents of Microbial2025 and compares them to an earlier database, Microbial2023, which was constructed following an approach described previously [10] and which has been used in prior metagenomic studies [11]. In addition, both databases include two human reference genomes (CHM13 and GRCh38) along with common laboratory vector contaminants from UniVec and EmVec, allowing reads derived from humans or vectors (both of which are very frequently observed in metagenomics experiments) to be identified and classified correctly.

#### Masking the Microbial2025 database

Having established *l* = 3 as the optimal unit size, we next sought to determine the entropy threshold *s* required to effectively mask k-mers that were either low-complexity or were shared across distant taxa, while minimizing the loss of informative sequence content. To identify this threshold, we examined the entropy distribution of 31-mers shared between the CHM13 human reference genome and fungal genomes in the Microbial2025 collection. K-mers shared between humans and fungi are generally not due to shared evolutionary history, but instead are enriched for low-complexity, repetitive elements that are prone to spurious cross-kingdom matches, making this shared set a useful basis for defining a threshold applicable across the broader database.

The resulting entropy distribution, shown in **Figure 2**, displayed a clear bimodal pattern, with a lower-entropy population centered around an entropy value of approximately 2.2, largely reflecting repetitive or low-complexity sequence, and a higher-entropy population centered around approximately 4.1, corresponding to more complex, potentially informative sequence shared between the two genome sets. To formally characterize this distribution, we fit a two-component Gaussian mixture model to the entropy values, allowing us to model each subpopulation independently and identify the point of minimal overlap between them.

**Figure 2.**
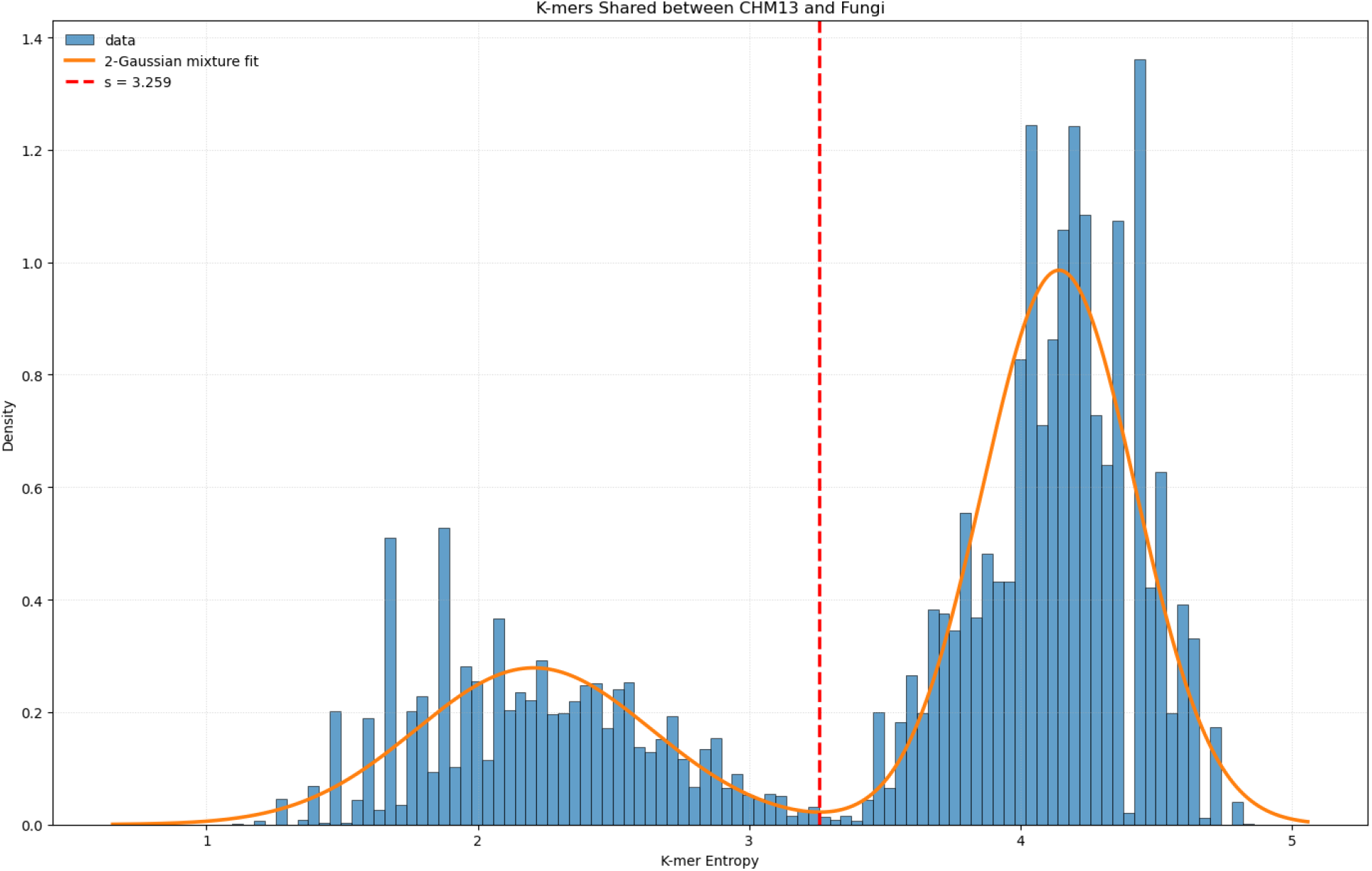
Entropy score distribution for 31-mers (unit size *l* = 3) shared between the CHM13 human reference genome and fungal genomes in the Microbial2025 collection. The bimodal distribution (blue bars) was fit with a two-component Gaussian mixture model (orange curve), revealing distinct low-entropy and high-entropy k-mer populations. The empirical entropy threshold (s = 3.259, dashed red line) was defined as the intersection point between the two fitted components and used to distinguish low-complexity k-mers (masked) from informative sequence content (retained) across the Microbial2025 database.

We then applied this same mixture-model-based algorithm to the set of Microbial2025 core genomes (human and vectors excluded), fitting a two-component Gaussian mixture model to the entropy distribution of all k-mers across this genome collection. This analysis identified an intersection point between the two Gaussian components at an entropy score of 3.30, which we adopted as the final entropy threshold. K-mers with entropy scores below this threshold were classified as low-complexity and masked, while those above the threshold were retained. This mixture-model-based approach provided a data-driven method for threshold selection, avoiding arbitrary cutoffs while ensuring the threshold reflected the natural separation between low- and high-complexity k-mer populations. The resulting threshold of *s* = 3.30 was applied uniformly across all genomes in Microbial2025.

### Masking reduced false classifications in TCGA data

To evaluate real-world performance, we began with a previously published set of 3,332,522 reads that all originated from low-count species in an analysis that classified thousands of human tumor samples from The Cancer Genome Atlas (TCGA) against the Microbial2023 database [7]. These classifications were flagged as suspect because only 1-10 reads from a particular species were identified in a given sample. Low-abundance species are commonly associated with contamination, cross-kingdom misclassification, or other sources of false-positive taxonomic assignment. We therefore reclassified this entire set of reads against the new Microbial2025 database to assess whether our masking had changed the original assignments.

Reclassification against Microbial2025 resulted in four outcome categories relative to the original Microbial2023 calls, summarized in **Figure 3**. The largest shift involved reads that were no longer classified, which means that the k-mers used to identify those reads were masked out in the newer database. This group contained 1,136,527 reads, 34.1% of the original set. Given that these reads were selected specifically as deriving from likely spurious matches, their elimination in this experiment suggests that they were false positives caused by low-complexity sequence, and demonstrates the effectiveness of the masking procedures applied during construction of Microbial2025.

**Figure 3.**
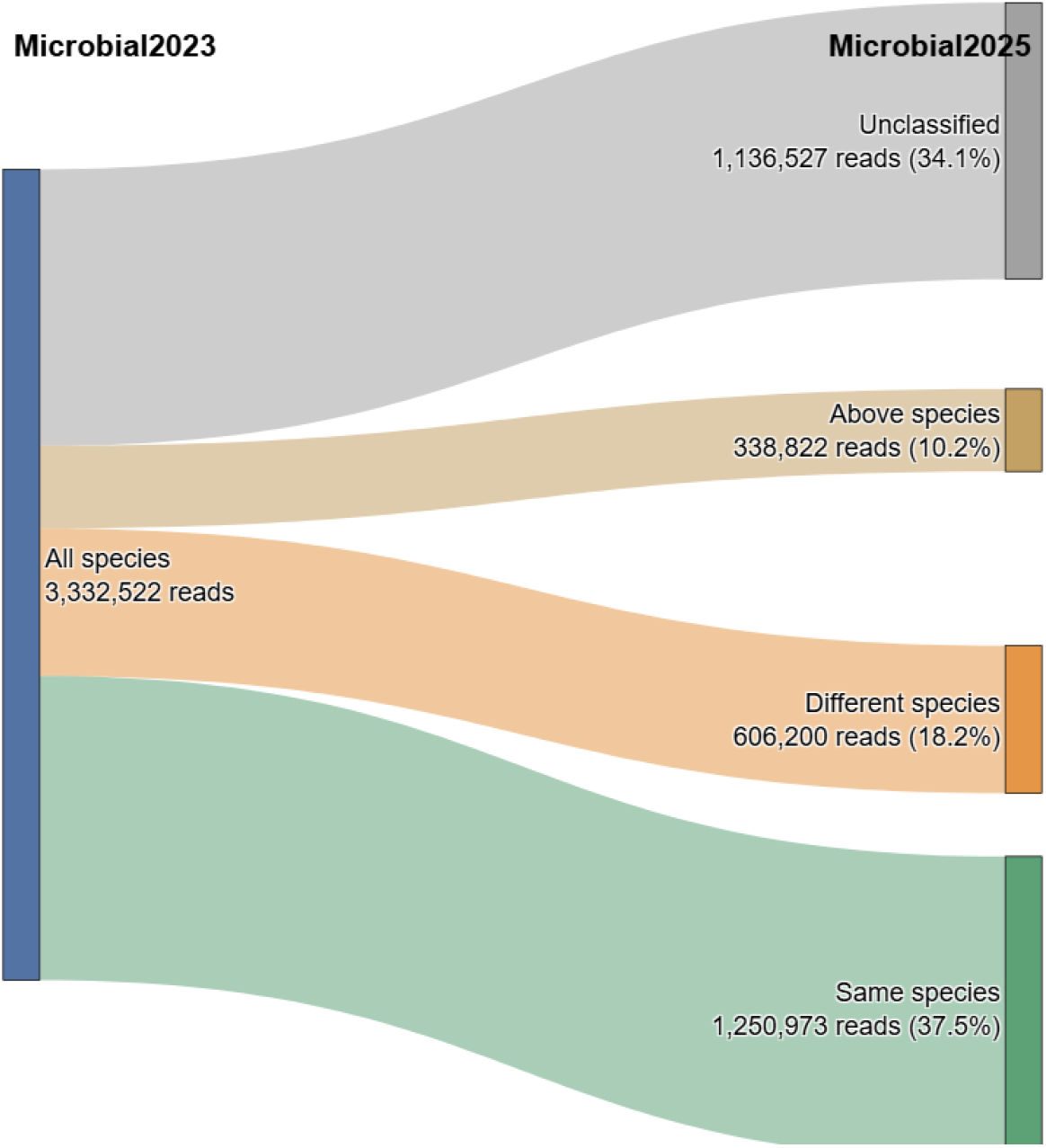
Sankey diagram tracking the reclassification of 3,332,522 suspicious, low-count reads from the TCGA cancer dataset, originally classified against Microbial2023, following reclassification against the masked Microbial2025 database. Reads are grouped into four outcome categories: unclassified under Microbial2025 (34.1%), indicating elimination of likely false-positive assignments; same species-level classification (37.5%); reassignment to a different species (18.2%); and resolution to a taxonomic rank above the species level (10.2%).

Among reads that remained classified under Microbial2025, 1,250,973 reads (37.5%) retained the same species-level assignment as under Microbial2023, indicating agreement between the two database versions for a substantial fraction of reads and supporting the retention of genuine, correctly classified signal. In contrast, 606,200 reads (18.2%) were reassigned to a different species, reflecting disagreement between the two versions, while the remaining 338,822 reads (10.2%) could only be resolved to a taxonomic rank above the species level (genus or higher), suggesting that Microbial2025 contains multiple species sharing the same 31-mer used for the earlier classification.

Taken together, these results indicate that Microbial2025 substantially reduces the number of low-count, likely false-positive classifications while preserving agreement with many other previous species-level assignments, supporting improved specificity in pathogen detection without a corresponding loss of true classification signal.

### Kmask achieves comparable specificity to SDUST with less masking

To directly compare Kmask against SDUST, an established and widely used low-complexity identification method, we applied both tools to the Microbial2025 collection (with human genomes excluded) and evaluated their effects on classification accuracy and masking efficiency. Using the same CHM13-derived 31-bp pseudo-read benchmark described above, Kmask achieved a false positive rate very similar to that of SDUST (5.78% vs. 5.17%), while masking substantially fewer total bases across the database (1.33% vs. 1.85%). This indicates that Kmask achieves comparable specificity in suppressing spurious human-microbial classifications while removing less sequence overall, representing a more efficient masking strategy per base removed.

To further characterize the differences between the two methods, we examined the length distribution of masked regions produced by each tool (**Figure 4A**). SDUST masked a large number of very short regions, with over 40% of masked intervals falling at or near the minimum detectable length, before transitioning to a broader distribution extending into much longer masked segments, in some cases spanning several kilobases. In contrast, Kmask’s masked region lengths were more narrowly distributed and consistently longer at the lower end, with no masking below 31 bp, which was the minimum length possible using the parameters in this study. This experiment shows that SDUST’s algorithm is able to (and does) flag very short segments as low-complexity, while Kmask’s parameters lead it to mask fewer, longer segments, which are intended to be more useful with k-mer-based classifiers such as Kraken and KrakenUniq.

**Figure 4.**
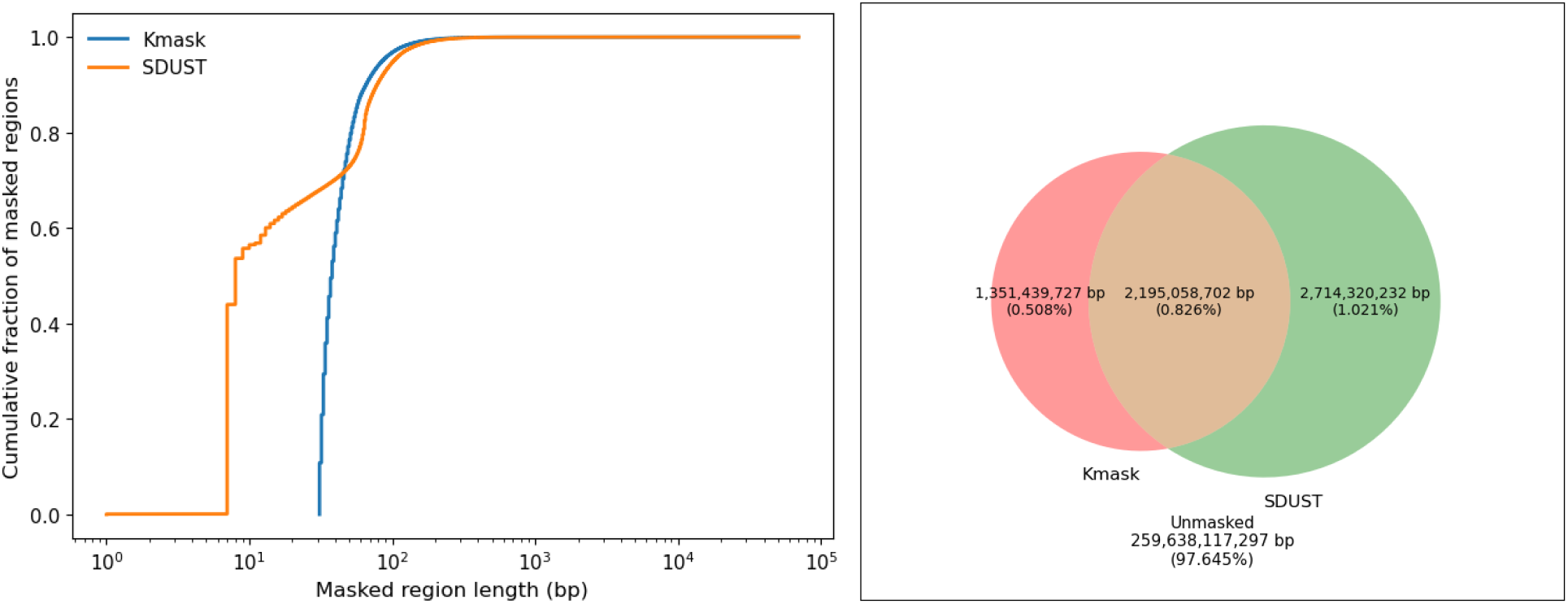
**A:** Empirical cumulative distribution function of masked region lengths produced by Kmask (blue) and SDUST (orange) across the Microbial2025 database. SDUST produced a substantial fraction of very short masked regions (7–10 bp), while Kmask masked regions were more narrowly distributed and consistently longer, reflecting its window-based entropy calculation and its 31-bp minimum masking length. **B:** Venn diagram showing the overlap in total bases masked by Kmask and SDUST across the Microbial2025 database. Of the database’s total bases, 0.508% were masked uniquely by Kmask, 1.021% uniquely by SDUST, and 0.826% by both methods, while the vast majority of bases (97.645%) remained unmasked by either tool.

We also assessed the overlap between bases masked by each method (**Figure 4B**). Interestingly, the two tools masked largely distinct sets of bases: while 2.2 Gbp were masked by both Kmask and SDUST, 1.35 Gbp were uniquely masked by Kmask, and 2.7 Gbp were uniquely masked by SDUST. The majority of the database (97.645%) remained unmasked by either method. This overlap indicates that Kmask and SDUST identify substantially different sequence features despite achieving similar overall reductions in false-positive classification, suggesting that the two approaches may be capturing complementary sets of low-complexity sequence.

## Methods

### Computation of Shannon entropy

To quantify sequence complexity, we computed the Shannon entropy across sliding windows of varying unit lengths. For a DNA sequence of length *k* and unit length of *l* (where *l* < *k*), we defined the Shannon entropy *H* as:

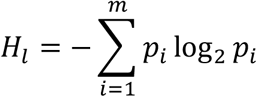

where *p*_*i*_ denotes the observed frequency of the *i*^th^ possible *l*-mer within the *k*-mer window, and *m* = 4^*l*^ represents the total number of possible *l*-mers. For example, *l* = 1 corresponds to mononucleotide entropy, whereas higher values of *l* (e.g., *l* = 2, 3, 4) capture di-, tri-, and tetra-nucleotide compositional complexity.

Each input genome sequence was scanned using a fixed-length sliding window aligned to Kraken’s *k*-mer length (*k* = 31 by default). For each *k*-mer, the per-window entropy score *H*_*l*_ was computed, and contiguous low-entropy windows below a user-defined threshold (*H*_*l*_ < *s*) were marked for masking.

**Figure S1.**
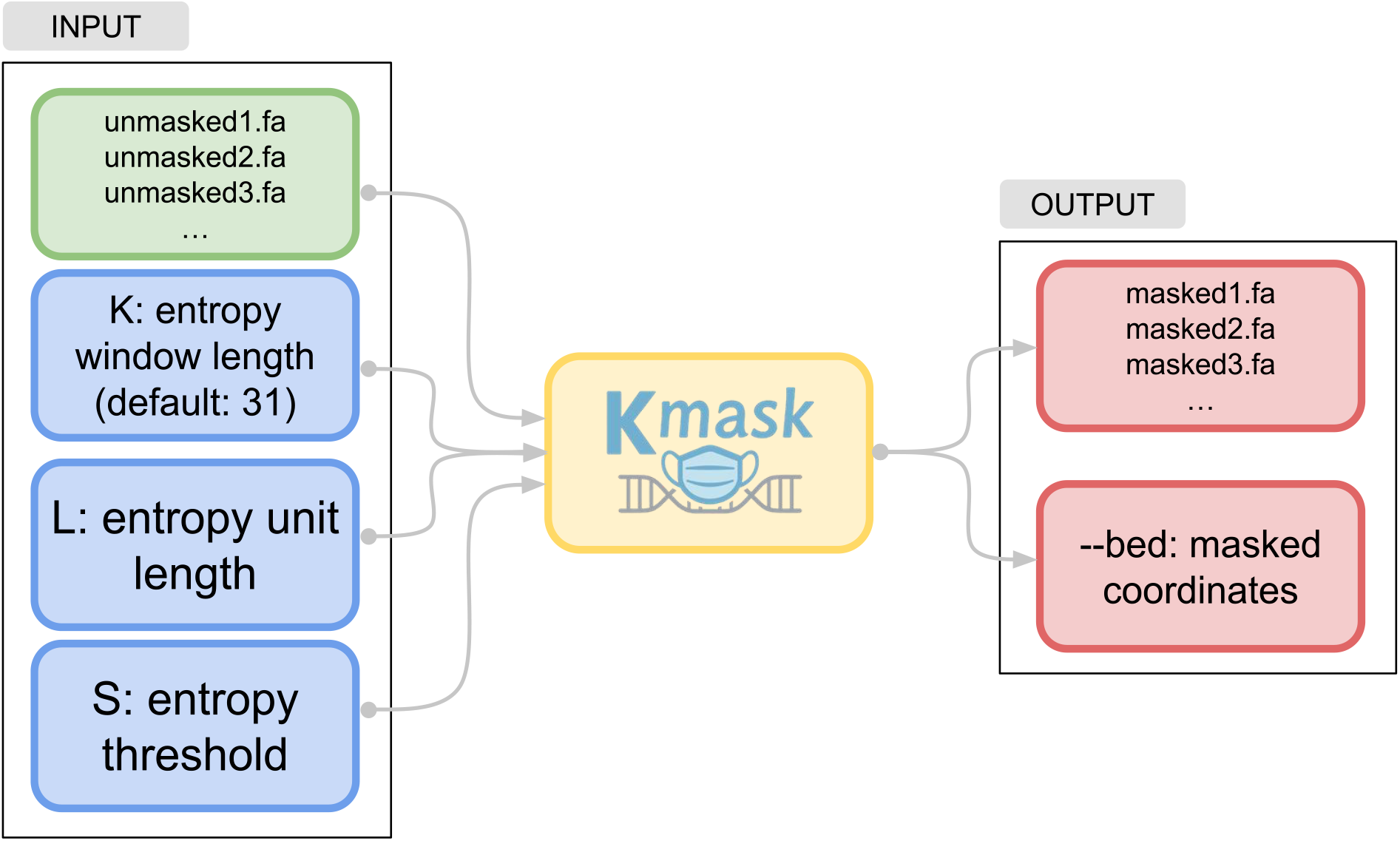
The Kmask processing flow. Kmask takes multiple FASTA files as input and processes them in parallel. It takes in three key parameters from the user: *k* for entropy window length (default: 31), *l* for entropy unit length (default: 3), and *s* for entropy threshold (default: 3.3). The outputs are a set of masked FASTA files, where low-entropy sequences are replaced by Ns, as well as a BED-format file specifying the coordinates of the masked regions.

### Efficient entropy computation via rolling updates

A naive implementation of sliding-window Shannon entropy would recompute the *l*-mer frequency distribution and entropy sum from scratch at every k-mer position, requiring *O(k)* work per window and *O*(*L* ⋅ *k*) total time across a genome of length *L*. To avoid this redundant computation, Kmask instead updates entropy incrementally as the window slides one base at a time, achieving overall *O(L)* time complexity independent of *k* or *l*.

We accomplish this using a circular queue (implemented as a fixed-size ring buffer) that tracks the *l*-mers currently contained within the active *k*-mer window. As the window advances by one position, exactly one *l*-mer exits the window (the “oldest” *l*-mer) and exactly one new *l*-mer enters (the “current” *l*-mer); all other *l*-mers within the window remain unchanged. The circular queue supports this update in constant time: rather than shifting all stored elements, the queue’s head and tail pointers are simply advanced modulo the queue’s fixed length, allowing the oldest *l*-mer to be evicted and the newest *l*-mer to be appended in constant time. See Supplementary **Algorithm S1** for a pseudocode description of this algorithm.

Because only two *l*-mer counts change between consecutive windows, and Shannon entropy is a sum over independent per-*l*-mer terms (*p*_*i*_log_2_*p*_*i*_), the entropy for the new window can be obtained by subtracting the two stale terms (corresponding to the old counts of the exiting and entering *l*-mers) and adding back the two updated terms (corresponding to their new counts), without recomputing the entropy contribution of any *l*-mer whose count did not change. This reduces the per-window entropy update from *O*(*k* ™ *l* + 1) (recomputing entropy over all constituent *l*-mers) to *O*(1), because only a constant number of frequency counts and logarithm evaluations are performed at each step, regardless of window size.

Together, the circular queue’s constant-time enqueue/dequeue operations and the incremental entropy update reduce Kmask’s total runtime from *O*(*L* ⋅ *k*) under a naive recomputation strategy to *O*(*L*), where *L* is genome length. This near-linear scaling is very helpful when applying Kmask to large reference collections such as Microbial2025, where entropy must be computed across tens of thousands of genomes comprising billions of bases.

### Selection of GC-diverse bacterial genomes

We selected twelve complete bacterial reference genomes from NCBI RefSeq [8] spanning a wide GC range (27–73%), including *Fusobacterium nucleatum* (low GC), *Escherichia coli* (medium GC), and *Frankia alni* (high GC). These genomes, shown in **Table 2**, were used to evaluate the effect of using different entropy thresholds for masking across genomes with different GC content. Each genome was processed using Kmask under a consistent set of masking parameters with *k*-mer length 31, *l*-mer unit size varying from 1–4, and entropy threshold *s* varying in a series of 21 linearly spaced values. For each combination of (*l, s*), all twelve bacterial genomes were masked, and the resulting masked genomes were used to construct a single KrakenUniq database. This approach yielded in total 84 control databases that differed only in the masking unit sizes (*l*) and thresholds (*s*) (see Supplementary **Table S1**). These experiments produced the ROC curves shown in **Figure 1**.

### K-mer-based screening of microbial assemblies against model organism references

To further test Kmask, we assembled an comprehensive microbial genome dataset spanning multiple taxonomic domains as an update to the early Microbial2023 database [11]. In total, we collected 71,210 genomes, including 55,247 bacteria, 651 archaea, 66 fungi, 14,583 viruses, and 663 eukaryotic pathogens. All genomes underwent a two-step cleaning pipeline to eliminate low-complexity regions and cross-species contaminants. This process ensures that the final database genomes contained only high-complexity, organism-specific sequences, reducing spurious alignments in downstream analyses.

First, all genomes were masked using both Kmask and SDUST. Windows with entropy below 3.3 were classified as low-complexity, and all bases in these regions were replaced with the N character. After this masking, each genome sequence consisted of only higher-complexity regions interspersed with N-masked gaps.

The next step was to remove high-complexity sequences that cause spurious mappings. Masking these sequences is important because many genome assemblies have been shown to be contaminated with small sequences from other unrelated species, a form of computational contamination [12,13] . In this step, all microbial genomes were screened against six model organism’s genomes plus a collection of widely-used laboratory vector sequences, shown in **Table 3**.

**Table 3:** Genomes used to screen the Microbial2025 database for accidental matches and contaminant sequences. EmVec and UniVec [14] are publicly-available collections of commonly used laboratory vectors and artificial sequences, which sometimes get incorporated by accident into newly assembled genomes.

| Species name | Taxonomy ID | RefSeq assembly ID |
| --- | --- | --- |
| *Homo sapiens* | 9606 | GCF_000001405.40 (GRCh38)<br>GCF_009914755.1 (T2T-CHM13v2.0) |
| *Mus musculus* | 10090 | GCF_000001635.27 (GRCm39) |
| *Drosophila melanogaster* | 7227 | GCF_000001215.4 (Release 6 plus ISO1 MT) |
| *Saccharomyces cerevisiae* | 559292 | GCF_000146045.2 (S288C) |
| *Danio rerio* | 7955 | GCF_000002035.6 (GRCz11) |
| *Caenorhabditis elegans* | 6239 | GCF_000002985.6 (WBcel235) |
| Synthetic construct | 32630 | EmVec & UniVec |

Each microbial genome was used as a query by KrakenUniq against databases created for each of the seven model organisms individually. KrakenUniq identified all exact 31-mer matches between the query and database genomes. When a segment of a microbial genome contained k-mers matching any of the model organism genomes (or synthetic constructs), all bases spanning that segment were replaced with Ns, as illustrated in **Figure 5**. By default, KrakenUniq classifies shared k-mers across taxa into their common ancestor, which can result in ambiguous classifications for k-mers conserved across multiple species. Classifying each genome against the seven contaminant species individually ensured that any match was attributed to a single contaminant genome.

**Figure 5:**
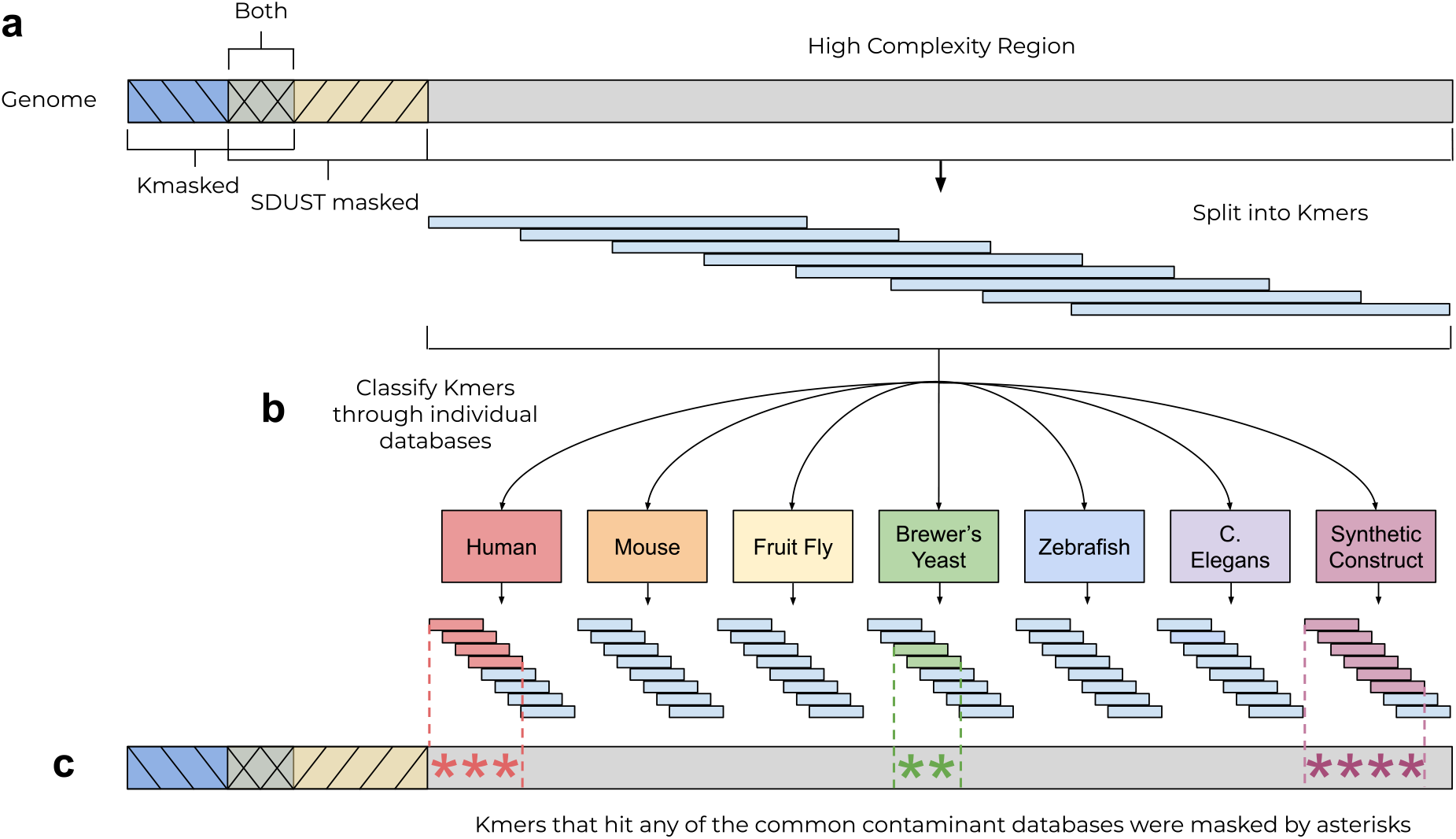
Shared K-mers filtering protocol. **a**) A microbial genome is first passed through Kmask and SDUST to mask out of its low-complexity sequences. **b**) The remaining high-complexity sequences are then compared to seven specialized KrakenUniq database. KrakenUniq internally splits the microbial genome into 31-mers and reports the locations of all 31-mers that have exact matches. **c**) Matching sequences are then masked out of the microbial genome.

We also did not mask certain matches that were deemed likely to reflect genuine homology rather than contamination. In particular, hits from the *S. cerevisiae* reference to fungal pathogen genomes were exempted from masking, because some fungal organisms share sequence similarity with *S. cerevisiae*. Similarly, matches to the synthetic construct database were ignored in bacterial or viral genomes, because some cloning vectors were designed using sequences from those microbes, so those overlaps were expected and were not treated as contamination.

## Discussion

We described Kmask, an entropy-based masking tool designed to address a persistent challenge in metagenomic classification: spurious taxonomic assignments arising from low-complexity sequence regions. By systematically optimizing two key parameters, we demonstrate that a carefully calibrated, information-theoretic approach to masking can substantially reduce false-positive classifications while preserving the informative sequence content needed for accurate pathogen detection.

When applied to the construction of Microbial2025, Kmask achieved a false positive rate only slightly higher than SDUST (5.78% vs. 5.17%) while masking substantially fewer bases (1.33% vs. 1.85%). This suggests that Kmask achieves a favorable trade-off between specificity and sequence retention, masking more efficiently per base removed. Unlike previous methods, Kmask prioritizes its masking strategy specifically for the k-mer length and matching logic that tools like KrakenUniq and Kraken2 use during classification. This distinction highlights that although Kmask can be used as a general-purpose low-complexity masking tool, its primary intended use is for k-mer-based metagenomic analyses. Also, because Kmask and SDUST mask different regions of a genome, we recommend using both tools when masking a database for use in either metagenomics or more general sequence alignment tasks.

Our validation using low-abundances species reported in a previous analysis of TCGA cancer samples further supports the utility of this approach. Reclassification against Microbial2025 eliminated more than a third of previously suspicious reads, indicating that the original matches were probably false positives caused by low-complexity matches. The reads that were reassigned to a different species or resolved only above the species level likely reflect a mixture of corrected misclassifications and cases where reduced k-mer availability (due to masking) limited resolution; distinguishing between these two possibilities represents an important direction for future validation.

Looking forward, Kmask’s near-linear time complexity and modular design make it well suited for integration into routine reference database construction pipelines beyond Microbial2025, including future updates to Kraken-family databases as new genome assemblies become available. More broadly, our results suggest that the benefits of low-complexity masking in metagenomic classification depend not only on the choice of complexity metric, but on the careful alignment of masking parameters with the alignment strategy of the downstream classifier itself, a consideration that may generalize to other k-mer-based bioinformatics tools beyond taxonomic classification, including genome sequence alignment and variant calling pipelines.

## Code and data availability

The Kmask software, including source code, installation instructions, and documentation, is freely available at https://github.com/yge15/kmask. The Microbial2023 and Microbial2025 KrakenUniq databases described in this study are freely available for download at https://benlangmead.github.io/aws-indexes/k2#krakenuniq.

## Acknowledgements

This work was supported in part by the US National Institutes of Health under grants R01-HG006677 and R35-GM130151.

## Supplementary Algorithm 1

Pseudocode for the entropy calculation in Kmask.

~~~
Input: A string s, positive integers k and l, where k > l.
Output: A sequence (s_i, e_i) of k-mers s_i extracted from s and their respective entropies e_i.
l_in_k <-k - l + 1
q <-{s[0:l), s[1:l+1), …, s[k - l:k)}
num_lmers = 4^l
lmer_counter <-{0, …, 0} // length num_lmers
for lmer in q do
    lmer_counter[lmer] += 1
done
cur_s <-s[0:k)
cur_sl <-s[k - l:k)
cur_e <-ShannonEntropy(lmer_counter)
output <-((cur_s, cur_e))
for i in [k:length(s)) do // rolling update of k-mers and Shannon entropy
    cur_s <-cur_s.pop_front().push_back(s[i]) // update k-mer
    cur_sl <-cur_sl.pop_front().push_back(s[i]) // update l-mer
    last_sl <-q.pop_front()
    q.push_back(cur_sl)
    if cur_sl != last_sl then
       last_se <-lmer_counter[last_sl] / num_lmers
       cur_se <-lmer_counter[cur_sl] / num_lmers
       lmer_counter[las  t_sl] <-lmer_counter[last_sl] - 1
       lmer_counter[cur_sl] <-lmer_counter[cur_sl] + 1
       new_last_sl = lmer_counter[last_sl] / num_lmers
       new_cur_sl = lmer_counter[cur_sl] / num_lmers
       // only calculate p*log2(p) for the entries that changed
         cur_e <-cur_e + last_se*log2(last_se) +
cur_se*log2(cur_se) - new_last_sl*log2(new_last_sl) -
new_cur_sl*log2(new_cur_sl)
      endif
      output.push_back((cur_s, cur_e))
done
return output
~~~

## Supplementary Table Captions

**Table S1. K-mask quality control datasets**.

**(a) Control genome panel**. Reference-quality RefSeq genome assemblies (n = 12) spanning a range of GC content (27.0–73.0%) and taxonomic groups, used as controls for k-mer masking validation. For each genome, the table lists organism name, strain, taxonomy ID, assembly name/accession, RefSeq annotation, assembly level, contig N50, genome size, submission date, gene count, and associated BioProject/BioSample identifiers.

**(b) CHM13 benchmarking**. Parameter sweep evaluating kmask performance against the CHM13 human reference assembly (n = 84 parameter combinations). For each combination of k-mer size (K), a secondary parameter (L), and masking stringency threshold (S), the table reports the number of k-mer hits, false positive rate (FPR), number of k-mers masked, database size, false negative rate (FNR), true positive rate (TPR), and area under the curve (AUC).

**Table S2. Reference genome assemblies used to construct Microbial2025 database**.

**(a–e) NCBI RefSeq/GenBank assembly summary records** for archaea (n = 651), bacteria (n = 55,247), fungi (n = 66), viruses (n = 14,583), and human (n = 2) genomes included in the k-mask reference database. Each record includes assembly accession, BioProject/BioSample identifiers, organism and strain information, assembly level and status, genome and annotation submitter, release date, genome size, GC content, replicon/scaffold/contig counts, and gene counts (total, protein-coding, and non-coding).

**(f) EuPathDB eukaryotic pathogen genomes** retrieved from EuPathDB release 68 (n = 663), listing file name, source organism, release version, category, file contents, file format, and file size, used to extend database coverage to eukaryotic pathogens.

